# Structure of a Single Chain H2A/H2B Dimer

**DOI:** 10.1101/2020.03.15.992685

**Authors:** Christopher Warren, Jeffrey B. Bonanno, Steven C. Almo, David Shechter

## Abstract

Chromatin is the complex assembly of nucleic acids and proteins that makes up the physiological form of the eukaryotic genome. The nucleosome is the fundamental repeating unit of chromatin, composed of ~147bp of DNA wrapped around a histone octamer formed by two copies of each core histone: H2A, H2B, H3 and H4. Prior to nucleosome assembly, and during histone eviction, histones are typically assembled into soluble H2A/H2B dimers and H3/H4 dimers and tetramers. A multitude of factors interact with soluble histone dimers and tetramers, including chaperones, importins, histone modifying enzymes, and chromatin remodeling enzymes. It is still unclear how many of these proteins recognize soluble histones; therefore, there is a need for new structural tools to study non-nucleosomal histones. Here we created a single-chain, tailless *Xenopus* H2A/H2B dimer by directly fusing the C-terminus of H2B to the N-terminus of H2A. We show that this construct (termed scH2BH2A) is readily expressed in bacteria and can be purified under non-denaturing conditions. A 1.31Å crystal structure of scH2BH2A shows that it adopts a conformation nearly identical to nucleosomal H2A/H2B. This new tool will facilitate future structural studies of a multitude of H2A/H2B-interacting proteins.

## 1. INTRODUCTION

The diploid human genome is composed of over six billion base pairs (bp) of DNA, which would be approximately two meters long if stretched end-to-end. This amount of genetic information needs to be condensed approximately 10,000-fold to be accommodated in the micron-scale cell nucleus. To achieve this level of compaction, eukaryotes wrap their DNA around highly basic histone proteins. The reversible, non-covalent interactions between DNA and histone proteins allow histones to be deposited on, slid along, or removed from DNA to enable essential cellular functions such as DNA replication, transcription and damage repair. The assemblies of DNA, histones and other associated proteins that make up the physiological form of the genome are collectively referred to as chromatin^1^.

Core histones (H2A, H2B, H3 and H4) are small, highly basic proteins that fold into a compact domain composed of three helices connected by short loops, with intrinsically disordered N- and C-terminal tails of variable length. The long central helix (α2) is flanked by two shorter helices (α1 and α3) connected by loops 1 (L1) and 2 (L2), respectively^2^. This histone fold mediates dimerization of complementary histones via a “handshake” interaction, where histones interact in an antiparallel fashion such that L1 of one histone is adjacent to L2 of the partner histone. The heterodimers are held together largely by hydrophobic interactions between the interacting histones.

In the nucleosome, the histone octamer organizes 147bp of DNA wrapped about the histones. The DNA is organized in ~1.7 negative superhelical turns with pseudo two-fold symmetry with each histone dimer pair organizing ~30bp of DNA^3,4^. Histone proteins make largely electrostatic contacts with DNA. Specifically, there are both positively charged amino acids and partial positive charges from helix dipoles interacting with the negatively charged phosphate groups on the DNA backbone, with very few interactions being made between the histones and bases of the DNA. Histone:histone contacts are made throughout the octameric structure, with 4-helix bundles formed between H3:H3 and H2B:H4. Nucleosomes are variably spaced, with linker DNA ranging from 20-80bp long. To facilitate compaction of chromatin fibers, linker histones, such as H1, are able to bind nucleosomes and linker DNA at the DNA entry and exit points^5–7^.

Histones are heavily post-translationally modified, mostly on their tails by methylation, acetylation, phosphorylation, ubiquitination, and sumolyation. These chemical modifications, along with modifications to the DNA itself, are “written” and “erased” by various enzymes and can recruit different “reader” proteins, which in turn can recruit other regulators of gene expression. These chemical modifications are heritable, as they can be propagated through cell division and possibly across generations in eukaryotic organisms^8–10^. While writer, eraser, and reader proteins are central for establishing and maintaining epigenetic states, other proteins known as histone chaperones and ATP-dependent chromatin remodelers are necessary to deposit and position histones on DNA^11^.

Histone proteins, therefore, have a large interaction network both on and off DNA. While there are many structures of nucleosomes and nucleosome-containing protein complexes, there are far fewer reported structures of individual histone dimers and tetramers, and their associated complexes. Many of the structures of proteins bound to H2A/H2B dimers have utilized either tailless or covalently linked constructs^12–16^. These constructs are likely necessary to stabilize the non-nucleosomal H2A/H2B dimer and limit flexibility for crystallography or NMR analysis. Here we describe a new single-chain construct derived from the *Xenopus laevis* H2A/H2B dimer, in which the histone tails are truncated and the C-terminus of H2B is directly fused to the N-terminus of H2A without an artificial linker sequence. This construct, termed single-chain H2BH2A (scH2BH2A), is readily expressed in bacteria, easily purified under non-denaturing conditions, and reproducibly yields crystals that diffract to high resolution. The structure of scH2BH2A shows that it folds into a nearly identical conformation as that observed in nucleosomal H2A/H2B, indicating that this tool may be useful in future structural studies of many H2A/H2B binding proteins.

## 2. METHODS

### Cloning, Expression and Purification of scH2BH2A

Overlapping PCR methods were used to remove the N-terminal tail of *Xenopus* H2B and directly fuse the C-terminus of H2B to the N-terminal helix of *Xenopus* H2A without an artificial linker sequence, while also removing the C-terminal tail of H2A (H2B residues 34-126 fused to H2A residues 14-105). This PCR product was cloned into the pRUTH5 vector containing an N-terminal His6 tag followed by a TEV protease site and glycine residue (His6-TEV-Gly-scH2BH2A). This construct was transformed into BL21(DE3) Rosetta2 cells and grown in LB media to an OD600 = 0.7. Protein expression was induced by adding 1mM IPTG to the culture and shaking for 3 hours at 37°C. Cells were pelleted and lysed by sonication in lysis buffer (25mM Tris pH 8.0, 1M NaCl, 1mM EDTA, 1mM PMSF, and 5mM BME). Lysate was clarified by centrifugation at 14,000rpm and supernatant was incubated with Ni-NTA resin while rotating for 1 hour at 4°C then passed over a gravity column. The resin was extensively washed with lysis buffer including 10mM imidazole and eluted with lysis buffer including 360mM imidazole. Imidazole was removed by dialysis into lysis buffer and the His6-tag was cleaved by overnight incubation with TEV protease. Subtractive Ni-NTA chromatography was used to remove the cleaved tag, any uncleaved target protein, and His6-tagged TEV. Flow through was concentrated and purified by size-exclusion chromatography over a Superdex 75 column equilibrated in 25mM Tris pH 8.0, 1M NaCl, and 1mM EDTA. scH2BH2A eluted from the column as a single peak and was >95% pure by SDS-PAGE analysis (Figure 1A). scH2BH2A was aliquoted, flash frozen and stored at −80°C.

**Figure 1 –.**
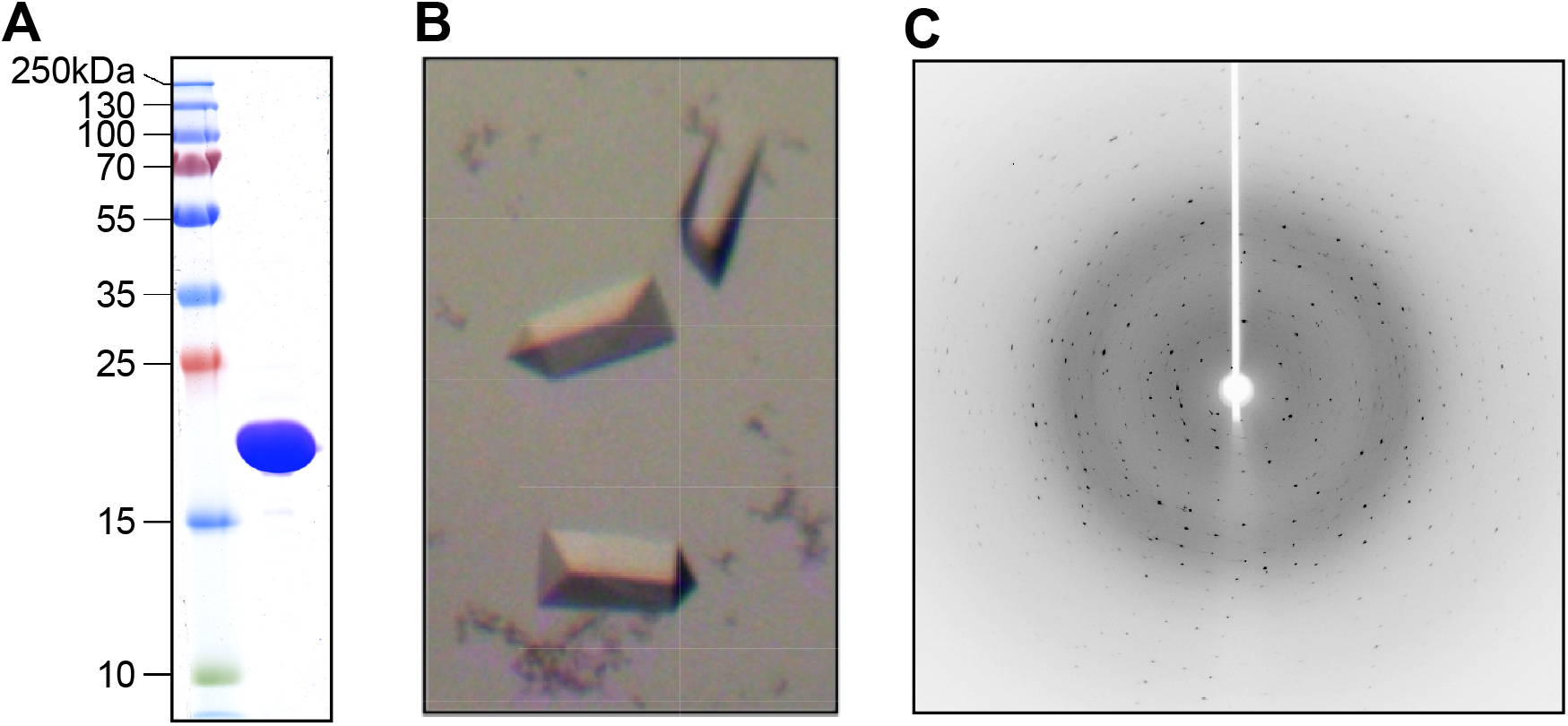
Purification and Crystallization of scH2BH2A. (A) 0.5μL of purified and concentrated scH2BH2A was run on 15% SDS-PAGE and stained in Coomassie Brilliant Blue. (B) Representative crystals of scH2BH2A. (C) Representative X-ray diffraction of a scH2BH2A crystal.

### Crystallization of scH2BH2A

scH2BH2A precipitated at lower salt concentrations. Based on buffer conditions used in another crystal structure of a peptide from the Spt16 subunit of FACT bound to H2A/H2B, we reasoned that inclusion of a non-detergent sulfobetaine (NDSB) in the buffer might increase solubility at low ionic strength^17^. NDSBs are commonly used as additives in protein crystallization screens for their ability to stabilize proteins and prevent aggregation^18^. An aliquot of scH2BH2A was thawed and rapidly diluted 20-fold in 25mM Tris pH 7.5, and 600mM NDSB-256 (Hampton), to give a final buffer composition of 25mM Tris pH 7.5, 50mM NaCl, 0.05mM EDTA, 570mM NDSB-256. scH2BH2A was concentrated to 18 mg/mL in a 10KDa MWCO Amicon centrifugal filter unit. scH2BH2A was screened for crystallization conditions using the Microlytic MCSG suite by sitting drop vapor diffusion using 0.3μL of protein and 0.3μL of well solution. An initial hit from MCSG-1 drop D4 produced a single crystal that was not suitable for structure determination. This condition contained 0.2M sodium thiocyanate pH 6.9 and 20% PEG3350. We expanded on this initial hit using the Hampton Silver Bullets additive screen. Diffraction quality crystals grew in 48 hours in hanging drop format using 1μL of protein and 1μL of well solution in drop F1 (Figure 1B), which contained the original buffer and precipitant with additives methylenediphosphonic acid, phytic acid, sodium pyrophosphate, and sodium triphosphate.

### Data Collection

Crystals were cryo-protected in well solution supplemented with 20% glycerol and flash frozen in liquid N2. scH2BH2A crystals were screened at the Advanced Photon Source (APS) LRL-CAT beamline 31-ID-D at a wavelength of 0.97931Å (Table 1, Figure 1C).

**Table 1:**
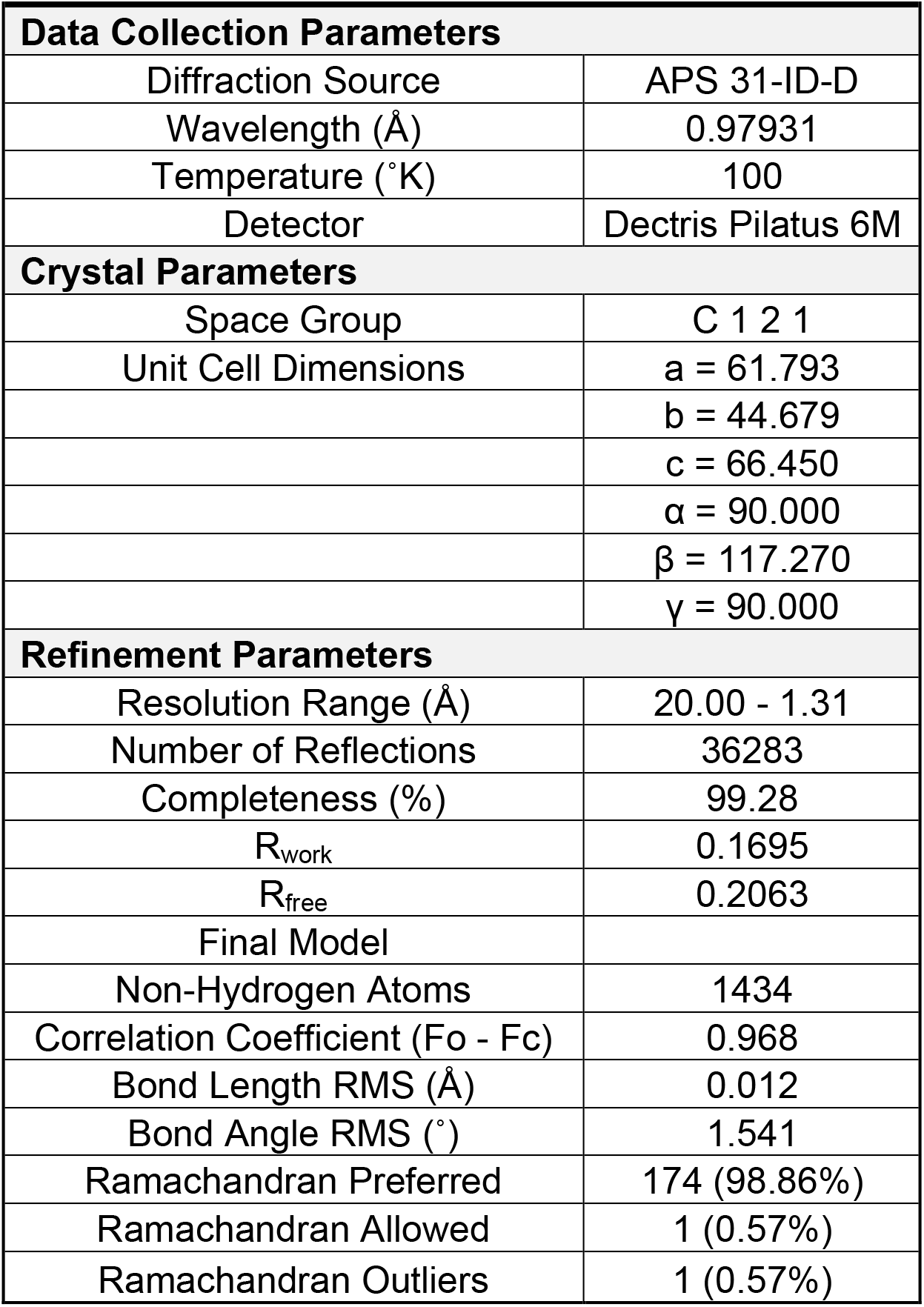
Data Collection and Structure Refinement

| **Table 1: Data Collection and Structure Refinement** |  |
| --- | --- |
| **Data Collection Parameters** |  |
| Diffraction Source | APS 31-ID-D |
| Wavelength (Å) | 0.97931 |
| Temperature (°K) | 100 |
| Detector | Dectris Pilatus 6M |
| **Crystal Parameters** |  |
| Space Group | C 1 2 1 |
| Unit Cell Dimensions | a = 61.793 |
|  | b = 44.679 |
|  | c = 66.450 |
| | $\alpha$ = 90.000 |
| | $\beta$ = 117.270 |
| | $\gamma$ = 90.000 |
| **Refinement Parameters** |  |
| Resolution Range (Å) | 20.00 - 1.31 |
| Number of Reflections | 36283 |
| Completeness (%) | 99.28 |
| $R_{\text{work}}$ | 0.1695 |
| $R_{\text{free}}$ | 0.2063 |
| Final Model |  |
| Non-Hydrogen Atoms | 1434 |
| Correlation Coefficient (Fo - Fc) | 0.968 |
| Bond Length RMS (Å) | 0.012 |
| Bond Angle RMS (°) | 1.541 |
| Ramachandran Preferred | 174 (98.86%) |
| Ramachandran Allowed | 1 (0.57%) |
| Ramachandran Outliers | 1 (0.57%) |

### Structure Solution and Refinement

Diffraction data were processed with HKL3000 software, and molecular replacement phasing, building and refinement were performed using the CCP4 suite and Coot software^19–21^. Diffraction extended to a resolution of 1.31Å and was consistent with the monoclinic space group C2, with cell dimensions a=61.793, b=44.679, c=66.450, α=90.000, β=117.270, γ=90.000 and one molecule of scH2BH2A per asymmetric unit. The structure was determined by molecular replacement using chains C and D (H2A and H2B) from the crystal structure of the *Xenopus laevis* nucleosome with histone tails truncated (PDB 1KX5)^3^. Iterative cycles of building and refinement procedures, including anisotropic B-factor refinement in later stages^22^ yielded a final model including 176 residues in a continuous chain consisting of H2B residues 36 to 126 followed by H2A residues 14 to 98 (numbering according to Uniprot entries Q92130 and P06897; the N-terminal cloning artifact glycine and H2B residues 34 and 35 along with H2A C-terminal residues 99 to 105 were not observed presumably due to disorder), 43 waters of solvation, and one molecule of pyrophosphate converging with Rwork and Rfree of 0.1695 and 0.2063, respectively (Table 1) and excellent stereochemistry. The coordinates were deposited in the PDB under the accession code 6W4L. All figures and structure comparison to nucleosomal H2A/H2B were performed with PyMOL software^23^.

## 3. RESULTS AND DISCUSSION

Purified scH2BH2A yielded reproducible crystals that consistently diffracted to high-resolution (Figure 1A-C). The 1.31Å structure of scH2BH2A shows that it forms the assembly expected for a H2A/H2B dimer (Figure 2A and B). The resolution of this structure allows unambiguous assignment of nearly all side chains along with 43 crystallographic water molecules (Figure 2C). A density feature not attributable to protein or water was modeled as a pyrophosphate molecule and was presumably derived from the additive crystallization screen condition. The pyrophosphate makes hydrogen bonding contacts with the backbone NH of Asp52 of H2B, and the side chain OH of next amino acid in the chain, Thr53. This pyrophosphate also makes multiple ionic interactions with the side chain of Arg72 of H2A in the same scH2BH2A molecule, as well as the side chain of K121 of H2B in a symmetry-related scH2BH2A molecule (Figure 2D). These contacts likely contribute to the observed propensity toward crystal formation.

**Figure 2 –.**
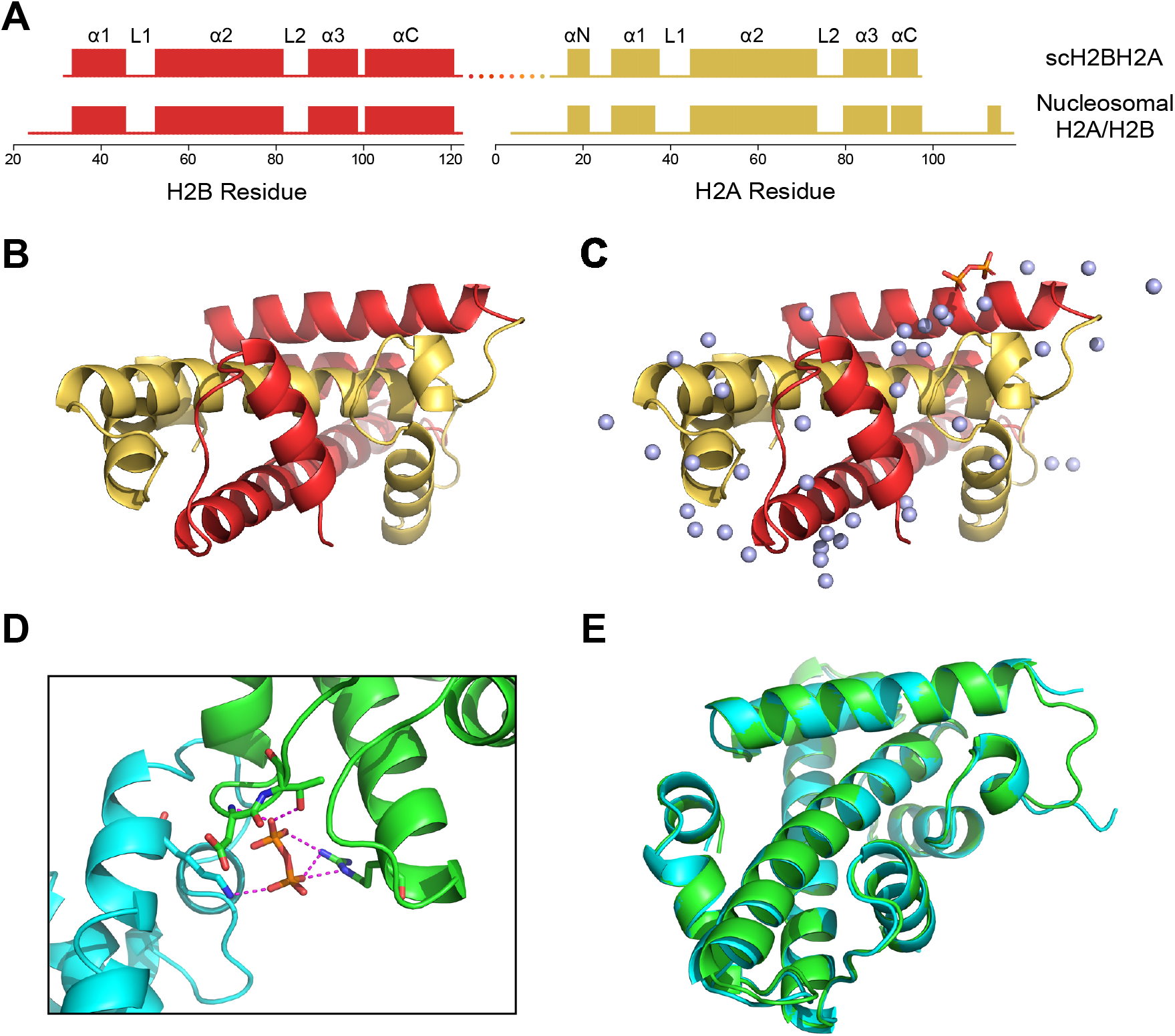
Structure of scH2BH2A. (A) STRIDE secondary structure alignment of scH2BH2A (top) with nucleosomal H2A/H2B (PDB 1AOI chains C and D, bottom). Conserved helices are shown as bars, the covalent link in scH2BH2A is shown as a dotted line. (B) Overall structure of scH2BH2A. Sequence corresponding to H2A colored yellow, sequence corresponding to H2B colored red. Linked histone tails are shown at the upper right portion of the structure. (C) Structure of scH2BH2A colored as in (B) with 43 crystallographic water molecules (blue spheres) and a single pyrophosphate molecule (orange and red sticks) also shown. (D) Structural details of scH2BH2A interactions with pyrophosphate at the crystallographic interface. Two symmetry related scH2BH2A chains are shown in green and blue. Electrostatic and H-bonding interactions between scH2BH2A and pyrophosphate shown as dashed lines. (E) Alignment of scH2BH2A structure with a nucleosomal H2A/H2B dimer. scH2BH2A colored green and nucleosomal H2A/H2B colored blue. Linked histone tails shown at the upper right portion of the structure.

Comparison of the scH2BH2A structure with nucleosomal H2A/H2B shows excellent agreement (Figure 2E). The positional RMSD between the two structures is 0.54Å based on 150 aligned C atoms. Each histone module in scH2BH2A exhibits the classic histone fold composed of three helices (α1, α2, α3) connected by two short loops (L1 and L2). Also apparent in the structure are the short N-terminal and C-terminal helices of H2A (αN and αC) and the longer αC of H2B. An extensive network of hydrophobic contacts contribute to the H2A/H2B interface. Notably, the αN helix of H2A is not apparent in a recent NMR solution structure of the human H2A/H2B dimer^24^. In the structure of the nucleosome, the H2A αN makes direct contacts with the phosphate backbone of DNA and is likely stabilized by these contacts. The presence of this short helix in the structure of scH2BH2A likely indicates that it is stabilized by the adjacent linked histone tails.

Finally, many attempts were made to co-crystallize and soak-in various short, acidic peptides derived from the H2A/H2B chaperone Nucleoplasmin (Npm2). We previously demonstrated that these peptides bind to H2A/H2B by tryptophan fluorescence and NMR^25^. However, no additional electron density was observed that could be assigned to these peptides in data arising from co-crystallization experiments, and crystals often cracked and dissolved during soaking experiments. We suspect that the tight packing and low solvent content of this crystal form makes introducing peptides difficult. It may be possible to introduce peptides that bind to a different surface of scH2BH2A; however, *de novo* co-crystallization trials are more likely to afford appropriate crystals of scH2BH2A bound to other protein and peptide ligands.

## Funding Sources and Acknowledgments

This work was supported by NIH R01GM108646, R01GM135614, and the American Lung Association (LCD-564723) (to D.S.), along with NIH F31GM116536 (to C.W.). The Albert Einstein Crystallographic Core X-Ray diffraction facility is supported by NIH Shared Instrumentation Grant S10 OD020068. Data collection also utilized resources of the Advanced Photon Source, a U.S. Department of Energy (DOE) Office of Science User Facility operated for the DOE Office of Science by Argonne National Laboratory under Contract No. DE-AC02-06CH11357. Use of the Lilly Research Laboratories Collaborative Access Team (LRL-CAT) beamline at Sector 31 of the Advanced Photon Source was provided by Eli Lilly Company, which operates the facility.

